# Integration of human and mouse single-cell transcriptomes of the developing cerebellum nominates cells-of-origin for Group 3 and 4 medulloblastoma

**DOI:** 10.64898/2026.01.22.701123

**Authors:** Ian Cheong, Leo Lau, Shraddha Pai

**Affiliations:** Ontario Institute for Cancer Research, Toronto, Canada; Department of Medical Biophysics, University of Toronto, Toronto Canada; University of Waterloo, Waterloo, Canada

## Abstract

The cerebellar rhombic lip neurogenic niche is critical for glutamatergic neurogenesis, and dysregulated differentiation of rhombic lip lineage cells is hypothesized to cause medulloblastoma (MB), a malignant pediatric cancer lacking targeted therapies. Humans have an expanded rhombic lip subventricular zone compartment not seen in mice or macaques, and the common Group 4 subtype of MB is hypothesized to arise by dysregulation of the EOMES+ unipolar brush cells (UBC) generated by this compartment. However, it is unknown whether humans have unique UBC populations not seen in the mouse and, if so, whether Group 4 MB tumour cells resemble these UBC populations. We integrated 336,598 human and mouse single-cell transcriptomes of the developing cerebellum (9–21 post-conception weeks in humans and E10–P14 in mice) and identified two subpopulations of UBC that are enriched in the human samples, relative to mice. One of these populations, which we term UBC 1, predominates at 11 post-conception weeks, shows upregulation of the Eyes Shut Homolog (*EYS*) transcript, is predicted to be driven by *OTX2*, *SOX4*, and *SOX11*, and is enriched for axonogenesis and neurodifferentiation pathways. We analyzed 27,735 single-cell transcriptomes from eight MB tumours and recapitulated the observation that Group 4 MB cells best resemble UBC. We then found that two-thirds of Group 3 and Group 4 MB UBC-like cells best resemble human-enriched UBC 1 cells. Gene regulatory network analysis revealed that top regulators of Group 4 MB tumour cells include *EOMES*, *SOX4*, and *SOX11*. Our work provides initial evidence for human-enriched UBC states, and suggests that *SOX4* and *SOX11* may drive neurogenesis in UBC and gene expression in a subset of Group 4 MB tumour cells. Our findings shed light on genetic determinants of human cerebellar expansion and suggest that human models may be needed to recapitulate the oncogenesis of Group 3 and 4 MB.

## Introduction

The cerebellum forms part of the hindbrain and is located below the cerebrum and behind the brain stem and fourth ventricle. Like the cerebrum, the cerebellum is divided into two hemispheres which are connected via the vermis, and it contains a cortical layer less than 1 mm thick in adults^1^. The cerebellar cortex itself is further divided into three layers: the molecular layer (outermost), the Purkinje cell layer, and the granule layer (innermost)^2^. Functionally, the cerebellum plays an important role in many day-to-day activities, including motor control, memory, speech and language, visuospatial processing, and executive function^3^.

Almost all that is known about cerebellar development comes from studies in mice. Initially, the cerebellar anlage is formed by the isthmic organizer through Fgf8 signalling around embryonic day (E) 8.5^4,5^. This is followed by the formation of the cerebellar ventricular zone (VZ) and the cerebellar rhombic lip (RL), the two main progenitor zones which give rise to all neurons of the cerebellum. The cerebellar VZ is specified by expression of *Ptf1a* and gives rise to all GABAergic neurons, including Purkinje cells and Pax2^+^ interneurons^4,5^. The Purkinje cells arise around E10.5–E13.5 and form the Purkinje cell layer of the cortex, with dendrites that also project into the molecular layer^2,4^. Meanwhile, the cerebellar RL is specified by expression of *Atoh1* and gives rise to all glutamatergic neurons such as granule cells (GCs) and unipolar brush cells (UBCs)^4,5^. The cerebellar RL first gives rise to the granule cell progenitors (GCPs) around E12.5, which migrate to form the external granule layer of the cerebellum^4,6^. Within the external granule layer, the postnatal proliferation and expansion of the GCPs is driven by Shh secreted from the Purkinje cells, before they finally migrate through the Purkinje cell layer and differentiate into GCs to form the (internal) granule layer^4,5^. RL production of UBCs, marked by expression of *Eomes*, occurs around E13.5. The UBCs first migrate into the developing white matter before finally settling in the granule layer around postnatal day P10^7^.

On the other hand, while the mouse provides a good starting point for understanding cerebellar development in humans, the process in humans is more complex and prolonged. Compared to mice, the surface area of the human cerebellum is 750-fold larger^5^. Furthermore, whereas the adult mouse cerebellum contains approximately 60% of all neurons in the brain, the adult human cerebellum contains approximately 80% of all neurons; this is equivalent to around 70 of the 86 billion neurons in the human brain^5,8^. Human cerebellar development begins at around 30 days post-conception and continues until around two years after birth^6^. Similar to mice, the human cerebellar VZ and RL give rise to GABAergic and glutamatergic neurons, respectively. Neurogenesis in the cerebellar VZ occurs until around 10 weeks post-conception (PCW), after which it begins to thin, and the cerebellar RL begins to expand. Several weeks later, the cerebellar RL embeds itself into the posterior-most lobule of the cerebellum where it persists until birth^6^. In contrast, the mouse cerebellar RL is much more transient and disappears by E17.5^6^. Additionally, the human cerebellar RL is elaborated into a rhombic lip ventricular zone (RL-VZ), which is KI67-rich and SOX2^+^, and a rhombic lip subventricular zone (RL-SVZ), which is KI67-rich and SOX2-sparse^6^. The RL-VZ and RL-SVZ are separated by a vascular bed, and progenitor cells in the RL-SVZ generate both the GC and UBC cell lineages^6^. This elaboration of the RL may be a feature unique to human neurodevelopment, as it is not seen in mice or even in rhesus macaques^6^.

Given the protracted and unique process of cerebellar development, especially in the cerebellar RL, humans may be particularly susceptible to insults causing various developmental diseases and disorders^5,9^. Medulloblastoma (MB) is hypothesized to result from failed differentiation during cerebellar development (see below). Neurodevelopmental disorders, such as some forms of autism spectrum disorder, attention deficit-hyperactivity disorder, and developmental dyslexia, are associated with abnormalities in the cerebellum^9,10^. Furthermore, genes that are implicated in Dandy-Walker malformation—a congenital disorder characterized by hypoplasia of the cerebellar vermis—and autism spectrum disorder show enriched expression across a number of cerebellar cell types^11,12^. As such, an improved understanding of the human-specific aspects of cerebellar development may provide valuable insights into the origins of these neurodevelopmental disorders.

MB is the most common malignant pediatric cancer of the central nervous system, accounting for approximately 20% of all childhood brain tumours and over 60% of all embryonal tumours^13,14^. It primarily affects children aged 0–9 at a rate of approximately 5 per million people, although cases do occur in adolescents and adults^13,14^. Overall, males are approximately 1.7 times more likely to be affected than females^14^. Treatment for MB generally involves with maximal safe resection of the tumour, followed by craniospinal irradiation and adjuvant chemotherapy with a combination of vincristine, cisplatin, cyclophosphamide, and lomustine^15,16^. Unfortunately, while patient survival may be as high as 80% post-treatment, survivors generally suffer a reduced quality of life and have an increased risk of hearing loss, reduced motor coordination, and cognitive and intellectual deficits^17–19^.

Molecularly, MB has been classified into four subgroups (WNT, SHH, Group 3, and Group 4), with each subgroup exhibiting unique gene expression patterns, mutations, DNA methylation profiles, and prognoses^15,20^. WNT and SHH MB are so named because they are driven by aberrant activation of the WNT and Sonic Hedgehog signalling pathways, respectively^20^. WNT MB makes up only around 10% of cases and has the best prognosis of all subgroups, with 5-year survival reaching up to 95%^21^. Over 85% of WNT MB patients have a mutation in the gene coding for β-catenin, *CTNNB1*^22^. The SHH subgroup encompasses approximately 30% of all MB cases, and prognosis is generally worse than that of the WNT subgroup^21^. Genes involved in the involved in the SHH pathway are frequently mutated (*PTCH1*, *SUFU*, *SMO*) or amplified (*GLI2*)^22^. SHH tumours have been shown to be transcriptomically most similar to GCP lineage cells, and these cells are hypothesized to be the cell of origin for this subgroup^23^.

Meanwhile, despite the fact that the Group 3 and Group 4 MB subgroups account for over 60% of cases, their molecular drivers remain poorly defined^21^. Preclinical models are severely lacking for these subgroups, especially for Group 4 MB, which has no cell lines^15^. A better understanding of the dysregulated genes driving Group 3 and Group 4 MB may help with the generation of useful models^15,16^. Recent studies have found that the transcriptomes of Group 3 and Group 4 MB tumours closely mirror those of the UBC lineage and its progenitors^23,24^. In particular, these tumours resemble UBCs and, notably, cells of the RL-SVZ^6,24,25^. Alterations in the core binding factor alpha (CBFA) complex have been found in Group 3 and Group 4 MB tumours; two of the CBFA complex members, *CBFA2T2* and *CBFA2T3*, are specifically expressed in the RL-SVZ^24^. Additionally, expression of *CBFA2T2* is inhibited by the transcription factor OTX2, and the *OTX2* gene is highly expressed in Group 3 and Group 4 MB^24^. Knockdown of *OTX2* in Group 3 MB cell lines, a proxy for Group 4 MB which lacks preclinical models, resulted in an upregulation of differentiation markers and *CBFA2T2* expression, suggesting that *OTX2* overexpression or loss of function of the CBFA complex could prevent differentiation and drive tumour formation^24,25^. Differentiation of SHH and Group 3 MB tumour cells can also be induced by thyroid hormone^26^, which is required for cerebellar development, further supporting the hypothesis that MB arises from dysregulated differentiation during cerebellar neurogenesis. Although the CBFA complex may be a key player in RL-SVZ differentiation and MB initiation, fewer than 60% of Group 4 MB tumours can be explained by mutations in the complex^24^, and more work is needed to identify other developmental programs promoting proliferation of these tumours.

Taken together, this suggests that the gene regulatory networks (GRNs) and molecular drivers of normal cerebellar development, when dysregulated, could be the same drivers of MB tumourigenesis; however, these developmental regulators are poorly understood. Furthermore, the fact that the RL-SVZ is a region of the cerebellum which is uniquely expanded in humans relative to mice leaves open the possibility that UBC cell states not readily apparent in the developing mouse cerebellum, but more abundant in the developing human cerebellum, may be the putative cells-of-origin for Group 3 and 4 MB. Therefore, understanding these GRNs could pave the way for identifying biomarkers unique to these human cell states or developing targeted therapies for tumour cells arising from these populations. We hypothesized that the UBC lineage of the developing human cerebellum contains cell states either missing or less abundant in the mouse, and that the GRNs from human-enriched cell states would resemble those in Group 4 MB cells. We therefore integrated single-cell transcriptomes of developing human and mouse cerebella to identify and characterize UBC cell states enriched in humans, relative to mice. We then analyzed single-cell transcriptomes from MB tumours to ascertain if Group 4 MB tumours resemble human-enriched cell states and what transcription factors these may have in common.

## Methods

### Datasets

This study involves secondary analysis of previously published data that is publicly available. Human samples had already been deidentified in these data. Similarly, this study only involves secondary analysis of mouse data, and no animals were used to generate new data for this work. Therefore, no ethics approval was obtained for this study for either human or animal research.

### Developing cerebellum

The counts matrices of two human and two mouse cerebellar single-cell RNA-sequencing (scRNA-seq) datasets^12,23,27^ were downloaded from the links in the published papers. Low quality cells or nuclei were filtered out as published. Based on the original cell type annotations of the datasets, the cell types were re-annotated and consolidated across datasets (Supplementary Tables 1-2).

### Medulloblastoma

MB scRNA-seq counts from Vladoiu *et al.*^23^ were kindly provided by Dr. Liam Hendrikse and Dr. Jiao Zhang (Dr. Michael Taylor Lab). The dataset contained 27,735 cells from eight MB tumours: two SHH tumours, two Group 3 MB tumours, and four Group 4 MB tumours. Counts from all cells were combined and normalized using *SCTransform*^28^. Batch correction was performed using fastMNN^29^. To visualize the cells, the top 50 MNN-corrected principal components (PCs) were used for dimensionality reduction with uniform manifold approximation and projection (UMAP).

### Integration of human and mouse datasets

To prepare the datasets for integration, the genes in the mouse datasets were unified with the human genes by identifying one-to-one orthologs (n = 16,755) using the *biomaRt* R package^30^. Each dataset was individually normalized using *SCTransform*^28^ with differences in the cell cycle phase regressed out. Dimensionality reduction was performed to visualize each dataset separately, as described in the next section.

Canonical correlation analysis (CCA) in Seurat^31^ was used to integrate the human and mouse cerebellar datasets (“full cerebellum integration”) (Supplementary Figure 1). To visualize the integration, dimensionality reduction was performed as described in the next section. Based on the consolidated cell types (Supplementary Table 2), the glutamatergic cells of the rhombic lip (RL) lineage (“RL”, “UBC/GCP progenitor”, “UBC”, “GCP”, and “GC”) and non-neuronal reference cell populations (“endothelial”, “microglia”, and “oligodendrocyte/OPC”) were isolated. Additionally, since expansion of the cerebellar RL occurs after 10 PCW^6^, human cells between 7 and 10 PCW were filtered out. Dimensionality reduction was performed on the isolated cells, and the cells were clustered using Louvain clustering (Seurat *FindClusters*, resolution = 0.4; “RL lineage integration” with 26,094 human cells and 55,449 mouse cells). The clusters were then annotated based on cell type marker expression and original cell type annotations (Supplementary Table 2).

The effect of CCA integration was compared to that with simple concatenation of the two datasets, and with integration using two additional approaches, reciprocal principal component analysis (RPCA) from Seurat^31^ and Harmony^32^ (Supplementary Figures 2–3). Overall, RPCA appeared to correct for species differences similar to CCA, whereas Harmony fully integrates cells from both species (Figure 1c, Supplementary Figure 3). However, when considering the originally annotated ventricular zone of the rhombic lip (RL-VZ) and subventricular zone of the rhombic lip (RL-SVZ) cells from the Aldinger dataset^12,24^, the RL-SVZ cells no longer cluster together in Harmony (Supplementary Figure 3e–f). On the other hand, both CCA and RPCA appear to better preserve the biology of the RL, with the majority of the RL-SVZ cells clustering together. All downstream analyses were based on CCA integration.

**Figure 1.**
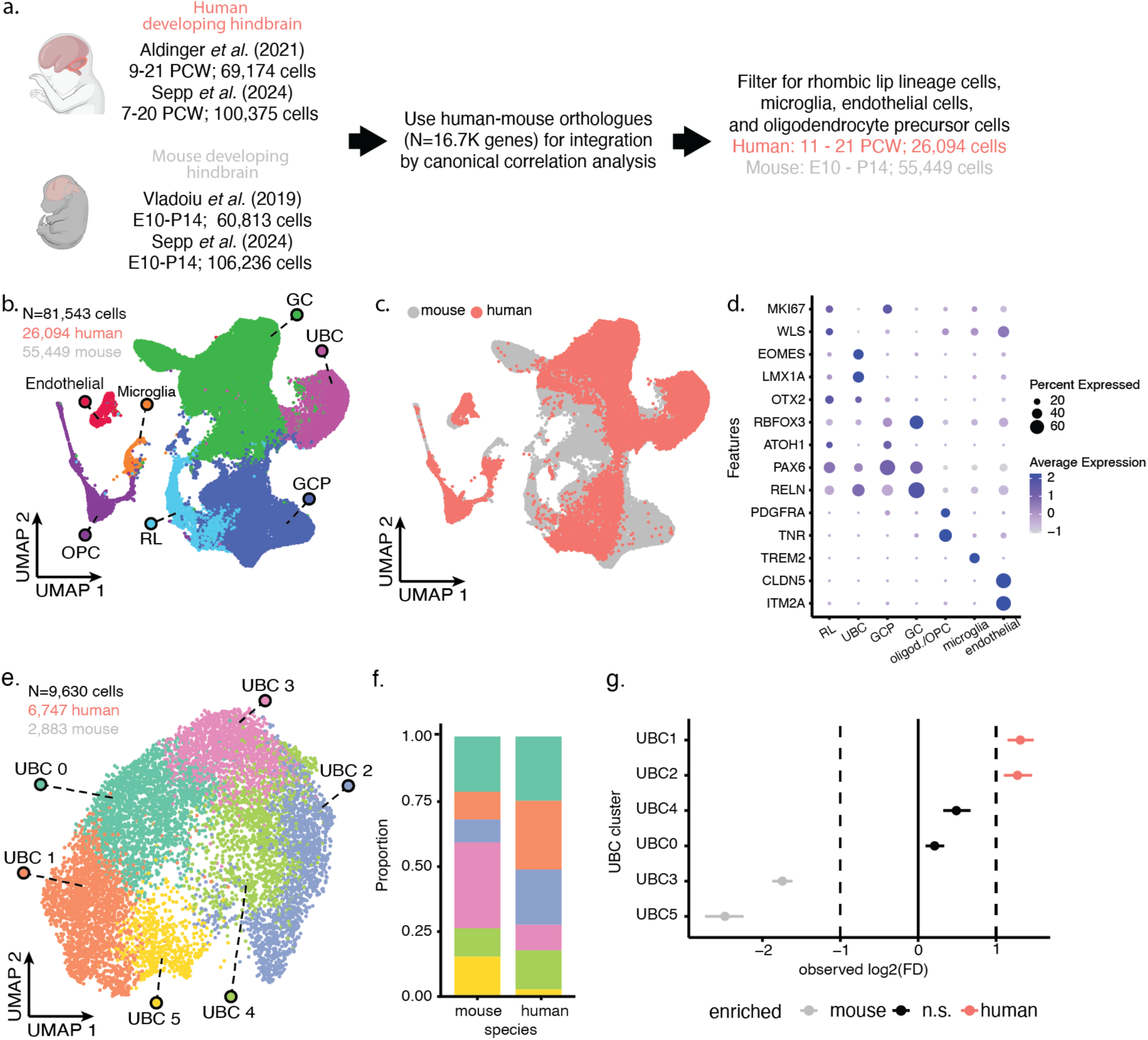
Integration of human and mouse UBCs reveal two UBC populations disproportionately present in humans. **a.** Schematic for human-mouse transcriptomic integration. **b–c.** UMAP visualization of integrated human-mouse cerebellar RL lineage cells and non-neuronal reference cells coloured by cell type (b) and species (c). **d.** Expression of marker genes for RL lineage cells and non-neuronal reference cells in the integrated data. **e.** UMAP visualization of subclustered UBCs. **f.** Proportion of human and mouse cells in each of the UBC clusters. **g.** Relative differences in proportion for each UBC cluster between humans and mice. Clusters with a log_2_FD > 1 are significantly enriched in humans, and vice versa for cell types with log_2_FD < −1 (FDR < 0.05, *n* = 10,000 permutations^34^).

To identify subclusters of unipolar brush cells (UBCs), the UBC cluster was isolated, and dimensionality reduction and Louvain clustering were performed. To determine the ideal number of subclusters, the UBCs were clustered at multiple resolutions ranging from 0.1 to 0.8 (Seurat *FindClusters*) and a clustering tree was generated to visualize the flow of cells between clusters across different resolutions (*clustree* package)^33^. Overclustering on the clustering tree appears as the formation of new clusters from multiple “parent” clusters, indicating instability in the clusters^33^. Given that a resolution of 0.4 showed signs of overclustering, a resolution of 0.3 (with six UBC subclusters) was used for downstream analyses (Supplementary Figure 4).

### Dimensionality reduction and visualization

Dimensionality reduction was performed using Seurat. In general, principal component analysis (PCA) was used to determine the top 100 PCs whenever dimensionality reduction was performed (*RunPCA*). The variance explained by each PC was plotted on an elbow plot, and based on the plot, the top 50 (full cerebellum integration) or 25 PCs were used for further dimensionality reduction with UMAP (*RunUMAP*).

### Testing for differences in cell type proportions

To test for differences in cell type proportions, the *scProportionTest*^34^ and *propeller*^35^ packages were used. Briefly, *scProportionTest* performs a permutation test (*n* = 10,000 permutations) to determine the 95% confidence interval for the log_2_FD where the FD (fold difference) is the ratio of the cell type proportion in humans to the cell type proportion in mice. Cell types with a |log_2_FD| > 1 and an FDR < 0.05 were considered to be enriched in one species or the other. Meanwhile, *propeller* uses sample-level proportions (biological replicates) to perform a moderated *t*-test using an empirical Bayes framework. As information about sample identity was not available for the Vladoiu dataset, cells were grouped by age, and cells from each age were considered to be a sample. Cell clusters with a Benjamini-Hochberg FDR < 0.05 were considered to be enriched in one species or the other.

### Differential gene expression and pathway enrichment analysis

Starting from the human and mouse RL lineage integration, we subset the human UBCs and identified differentially expressed genes in each of the UBC subclusters using Seurat. Prior to differential gene expression analysis, genes were first filtered to keep only those that were expressed in at least 1% of all UBCs in each of the human datasets, Aldinger and Sepp. 8,122 genes remained after filtering. Each dataset was then individually normalized using *SCTransform* with differences in the cell cycle phase being regressed out. The normalized datasets were combined and prepared for differential gene expression analysis (*PrepSCTFindMarkers*). Differentially expressed genes were identified by performing a Wilcoxon rank-sum test comparing cells from one UBC cluster to cells from all other UBC clusters combined; this analysis was performed separately for the Aldinger and Sepp datasets (*FindConservedMarkers*). Genes were only tested if they were expressed in at least 1% of cells in at least one of the tested groups (*min.pct = 0.01*). The resulting nominal *p*-values were corrected using the Bonferroni method, and genes with an adjusted *p*-value < 0.05 were considered significant. Differentially expressed transcription factors (TFs) were identified using the list of 1,639 human TFs from Lambert *et al.*^36^.

The *gprofiler2* R package, which provides access to the online g:Profiler tool^37,38^, was used to run a functional enrichment analysis (*gost*) on genes that were upregulated in both human datasets (log_2_FC > 0 and nominal p-value < 0.05 for both the Aldinger and Sepp human datasets). A custom Enrichment Map gene set was used (downloaded from https://download.baderlab.org/EM_Genesets/August_08_2023/Human/symbol/)^39^) which included pathways from GO Biological Processes (excluding those inferred from electronic annotation (IEA))^40^, HumanCyc^41^, MSigDB (hallmark, C2, and C3 collections)^42,43^, NCI-Nature^44^, NetPath^45^, Panther^46^, Pathbank^47^, Reactome^48^, and WikiPathways^49^. Only pathways with 10–250 genes were included (*n* = 8,404 pathways). The genes that were tested for differential expression were used as the background set for pathway analysis. Pathways with a Benjamini-Hochberg FDR < 0.05 were deemed significantly enriched. Pathway enrichment analysis of the human genes without a one-to-one mouse ortholog (“non-orthologous genes”) was performed in a similar manner. For this analysis, the non-orthologous genes were set as the query and all human genes from both human datasets, Aldinger^12^ and Sepp^27^, were used for the background.

### Projecting developmental signatures onto tumour cells with SingleR

The CCA-integrated human and mouse RL lineage cell dataset was used as the developmental reference for projection onto tumour cells after being subset to include only human cells (26,094 cells). Prior to using SingleR^50^ for the projection, gene expression in both the tumour and reference datasets were individually normalized using *NormalizeData* on the RNA slot. *SingleR* was then run with *de.method* set to “wilcox”, using the integrated cell type label for projections. Group 3 and Group 4 MB tumour cells that mapped to “UBC” cells (UBC-like tumour cells) were then subset for further projection (11,483 tumour cells). For the projection of the detailed UBC states, the reference dataset was subset to only include the human cells in the six UBC cell clusters uncovered in this work (UBC_i, i = {0, … 5}) (6,747 cells). This reduced human UBC-specific reference set was then projected onto the UBC-like tumour cells to ascertain similarity to the new UBC clusters.

### Inferring regulons with pySCENIC

Gene regulatory network inference was performed using pySCENIC^51^. Briefly, raw counts were used to infer co-expression modules of TFs and corresponding target genes using *GRNBoost2*. *cisTarget* was then used to predict regulons by searching for enriched TF-binding site motifs 10 kbp upstream and downstream of the transcription start site of the target genes. The activity of each regulon in each cell was then measured with *AUCell*, and the regulon specificity score (RSS) was calculated to identify the top regulons that are specific to each cluster/cell type.

### Inferring developmental stage with pseudotime

Pseudotime analysis was performed on the human RL cells, UBCs, granule cell progenitors (GCPs), and granule cells (GCs) using Monocle 3^52^. After subsetting the cells from the cross-species integration, PCA was performed to determine the top 100 PCs (*preprocess_cds*). UMAP was run on the top 25 PCs (*reduce_dimension*) with the minimum distance (*umap.min_dist*) set to 0.3 and the size of the local neighbourhood (*umap.n_neighbors*) set to 30. The trajectory of the cells was inferred (*learn_graph*) and the cells were ordered by pseudotime (*order_cells*) by setting the RL cell cluster as the starting point.

## Results

### Integration of human and mouse developing cerebellar single-cell transcriptomes reveals human enrichment of unipolar brush cells

We first processed two human and two mouse cerebellar datasets and integrated them using canonical correlation analysis (CCA) (N=336,598 cells in total; 169,549 human, 167,049 mouse; Figure 1a, Supplementary Table 1, Supplementary Figure 1a–d)^12,23,27,31^. Overall, the cells clustered together by cell type and not by dataset or species (Supplementary Figure 1e–g). In comparison, the unintegrated datasets showed clear dataset and species batch effects (Supplementary Figure 2). As we were primarily interested in the glutamatergic neurons that arise from the rhombic lip (RL), we isolated the RL lineage cells and three non-neuronal reference populations, and performed dimensionality reduction for visualization. As before, the human and mouse cells appeared to be well-integrated (Figure 1b–c, Supplementary Figure 5). We identified 18 clusters, with clusters generally containing cells from multiple datasets (Supplementary Figure 5d). The clusters were then annotated using known cell type markers (Figure 1b,d), such as *EOMES* and *LMX1A* for unipolar brush cells (UBC), *WLS* for RL cells, and *RBFOX3* for granule cells (GC) (Figure 1b,d, Supplementary Figure 6). The annotation of the clusters was verified using the original cell type annotation. We included non-neuronal cell populations of endothelial cells, microglia, and oligodendrocytes/oligodendrocyte precursor cells in the integration, to determine potential under-integration of human and mouse cells. Given that these non-neuronal cells from both species clustered together, it appears that CCA is appropriately correcting for the species batch effect (Figure 1b–c).

Comparing RL-lineage cell type proportions between mice and humans revealed an enrichment of UBCs in humans (Supplementary Figure 7). Across all the datasets, the UBCs make up 29% of the human RL-lineage cells compared to only 5.5% in mice. The enrichment of UBCs in humans was found to be significant by two different approaches (Q < 0.001, Supplementary Figure 7b-c)^34,35^. Meanwhile, the proportion of RL cells and granule cells (GCs) were not significantly different between the two species (log_2_FD < 1, Supplementary Figure 7b). Granule cell progenitors (GCP) appeared to be enriched in mice by some measures (Supplementary Figures 7b–c). This effect is likely seen because the data analyzed includes early postnatal timepoints in mice when GCP expansion occurs, but does not include human cells from the third trimester onwards, when a similar expansion occurs in the developing human cerebellum^5^.

### Enrichment of unipolar brush cell subpopulations

Given that the UBCs were enriched in humans compared to mice, we wondered if certain subclusters of UBCs were driving this enrichment. We subclustered UBCs and identified six UBC subpopulations, which we labelled UBC 0 to UBC 5 (Figure 1e). We found that the UBC 1 and UBC 2 populations are enriched in humans compared to mice (Figure 1f–g; Q < 0.001, log_2_FD > 1). In particular, the UBC 1 cells comprise 26.4% of human UBCs compared to 10.6% of mouse UBCs (Figure 1f). Similarly, the UBC 2 cells comprise 21.3% of human UBCs compared to 8.8% of mouse UBCs (Q < 0.001 and log_2_FD > 1). Using a second method that relies on per-sample proportions^35^, UBC 1 was not found to be significantly enriched, but UBC 2 was (UBC 1: Q > 0.1; UBC 2: Q < 0.025; Supplementary Figure 8). In humans, the majority of UBCs at 11 PCW appears to be UBC 1, whereas the proportion of UBC 2 cells increases and forms the majority of the human cells at 20 PCW, along with UBC 0 and UBC 3 (Supplementary Figure 9a). On the other hand, the earlier UBCs in mice are comprised of UBC 0 and UBC 5 cells, and the UBC 2 and UBC 3 cell populations arise peri- and postnatally (Supplementary Figure 9a). The UBC subpopulations do not appear to be driven by any single dataset; rather, they seem to be well-represented in both datasets (Supplementary Figure 9c). Similarly, the distribution of UBC subpopulations by sex appears to reflect the sex distribution of cells by dataset (Supplementary Figure 10) and does not appear to be driven by any one sex. Considering only UBC cells from the Aldinger dataset, which are more evenly balanced in terms of sex as compared to the Sepp dataset (Aldinger: 2,915 cells, 58.5% female; Sepp: 3,832 cells, 5.2% female), 60% of cells of the UBC 1 subpopulation is from female samples and 69.2% of the UBC 2 subpopulation is from female samples (Supplementary Figure 10).

### Characterization of human-enriched unipolar brush cells

To better understand the temporal distribution of the UBC 1 cells, we inferred the developmental trajectory and pseudo-temporal ordering of human cells in the RL lineage^52^. Consistent with previous analyses^4,12,23^, the RL trajectory splits into two to give rise to both GCs and UBCs (Figure 2a). The UBC 1 cells appear to arise directly after the RL cells, and the UBC 2 cells appear at the terminal end of the trajectory, consistent with the changes in proportion seen between 11 and 20 PCW (Figure 2b-c, Supplementary Figure 9).

**Figure 2.**
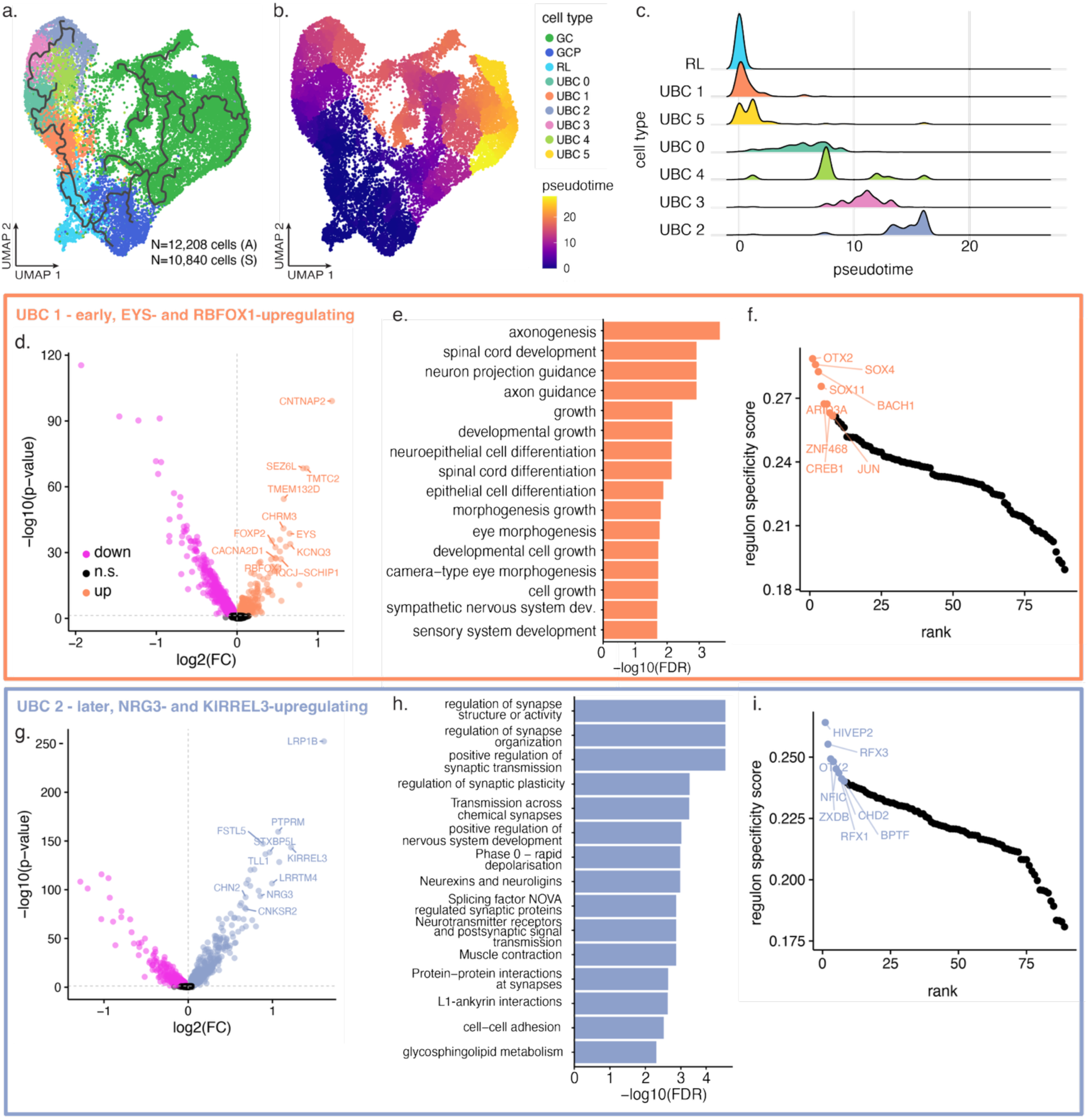
Characterization of human-enriched UBC cell states. **a–b.** UMAP visualization of the human cerebellar RL lineage cells coloured by cell type (a) and inferred pseudotime (b). The RL cells were set as the root cells and the inferred trajectory is overlaid in (a). **c.** Distribution of inferred pseudotime for cells of the unipolar brush cell lineage. **d.** Differentially expressed genes in the human-enriched UBC 1 cluster; top upregulated genes are labelled. Data shown for the Aldinger dataset. **e.** Top enriched pathways in genes upregulated in UBC 1 (Q < 0.05). **f.** Top regulons for UBC 1 cells. **g.** Differentially expressed genes in the UBC 2 cluster; top upregulated genes are labelled. Data shown for the Aldinger dataset. **h–i.** Enriched pathways (h) and top regulons (i) for UBC 2 cells.

A total of 73 genes were upregulated in the UBC 1 population of both Aldinger and Sepp datasets (7,928 genes tested; log_2_FC > 0 and Q < 0.05; Figure 2d, Supplementary Figure 11, Supplementary Table 3). This list included neurodevelopmental genes such as *CNTNAP2* and *FOXP2*, mutations in both of which have been linked to various intellectual and speech disorders^53,54^. Pathway enrichment analysis showed that genes upregulated in this cell cluster are enriched for pathways related to neuronal and axonal development (34 enriched pathways; FDR < 0.05, g:Profiler^55^; Figure 2e, Supplementary Table 4). We note that the UBC 1 cells also show an upregulated expression of Eyes Shut Homolog (*EYS*), a gene required for retinal photoreceptor cell function and loss-of-function mutations in which results in autosomal recessive retinitis pigmentosa, a retinal degeneration disorder^56^. Expression of a photoreceptor signature has previously been detected in the RL-SVZ and in a subset of Group 3 MB cells^25^. The *OTX2* gene regulatory network was found to be the most active in the UBC 1 cells (pySCENIC; Figure 2f). Other top regulons included those of *SOX4*, *SOX11*, *BACH1*, and *CREB1* (Figure 2f, Supplementary Table 5). *OTX2* has been shown to be overexpressed in MB and is hypothesized to maintain a less differentiated state in the RL^21,24^.

UBC 2 cells showed an upregulation of 459 genes, including *NRG3*, *CNKSR2*, and *KIRREL3*^57–59^, genes involved in neuronal differentiation and mutations in which are associated with neurodevelopmental disorders (Q < 0.05, log2FC > 0; Supplementary Table 3). Consistent with the fact that these cells arise later in development, genes upregulated in UBC 2 cells were enriched for pathways related to synaptic activity (Figure 2h; Supplementary Table 6). Top regulons enriched in UBC 2 cells included *OTX2*, but also *RFX3*, a transcription factor that serves as a master regulator of nervous system development and ciliogenesis, and disruption in which causes some types of autism^60–62^ (Figure 2i, Supplementary Table 5). Taken together, these results suggest that the UBC 1 cells represent a cell state that is relatively less differentiated than UBC 2.

### Shared drivers in unipolar brush cells and medulloblastoma

To ascertain if the human-enriched UBC states above are similar to Group 3 and 4 MB tumour cells, we analyzed published single-cell transcriptomes of 27,735 cells from SHH, Group 3, and Group 4 MB tumours (Figure 3a–b)^23^. We first determined the developmental best-matching cell state for each tumour cell, using human cells from our integrated dataset of cerebellar RL lineage cells and non-neuronal reference cells (Figure 3c–d). Nearly half of SHH tumour cells best resemble granule cell progenitors (42.4%); in contrast, < 2% of Group 3 MB (1.8%) or Group 4 MB cells do (1.1%). By comparison, over half of Group 3 MB tumour cells best resemble the RL progenitor cells (51.1%), and roughly one-third resemble UBCs (36.6%). Two-thirds of Group 4 MB tumour cells resemble UBC cells (69.8%), while most of the remaining cells resemble RL cells (27.4%) (Figure 3d). This profile of best-matching developmental cell states is consistent with the current model of developmental MB oncogenesis, which hypothesizes that SHH arise from dysregulated differentiation of GCPs, while Group 3 and 4 MB tumours arise from dysregulation of the RL, with Group 3 MB arising from a less differentiated cell state than Group 4 MB^23–25,63^. To further identify which of the six integrated UBC cell states Group 3 and 4 MB tumour cells best resemble, we projected signatures of the subclustered UBC populations onto UBC-like Group 3 and 4 tumour cells (11,483 tumour cells, 6,747 reference UBC cells). We found that two-thirds of Group 3 MB (68.6%) and Group 4 MB (69%) tumour cells best resemble the EYS-rich UBC 1 cell state, with the UBC 0 cell state accounting for roughly one-third of cells in each tumour type (24.7% Group 3 MB, 30.5% Group 4 MB cells; Figure 3e); this distribution does not appear attributable to any individual tumour sample (Supplementary Figure 12). While a small fraction of Group 3 and Group 4 MB cells mapped to the UBC 4 cell state (6.7% Group 3 MB, < 1% Group 4 MB), none of the cells mapped to the later, NRG3-rich UBC 2 cell state (Figure 3e). In summary, Group 3 and 4 MB cells that are UBC-like appear to best resemble the early UBC 1 cell state.

**Figure 3.**
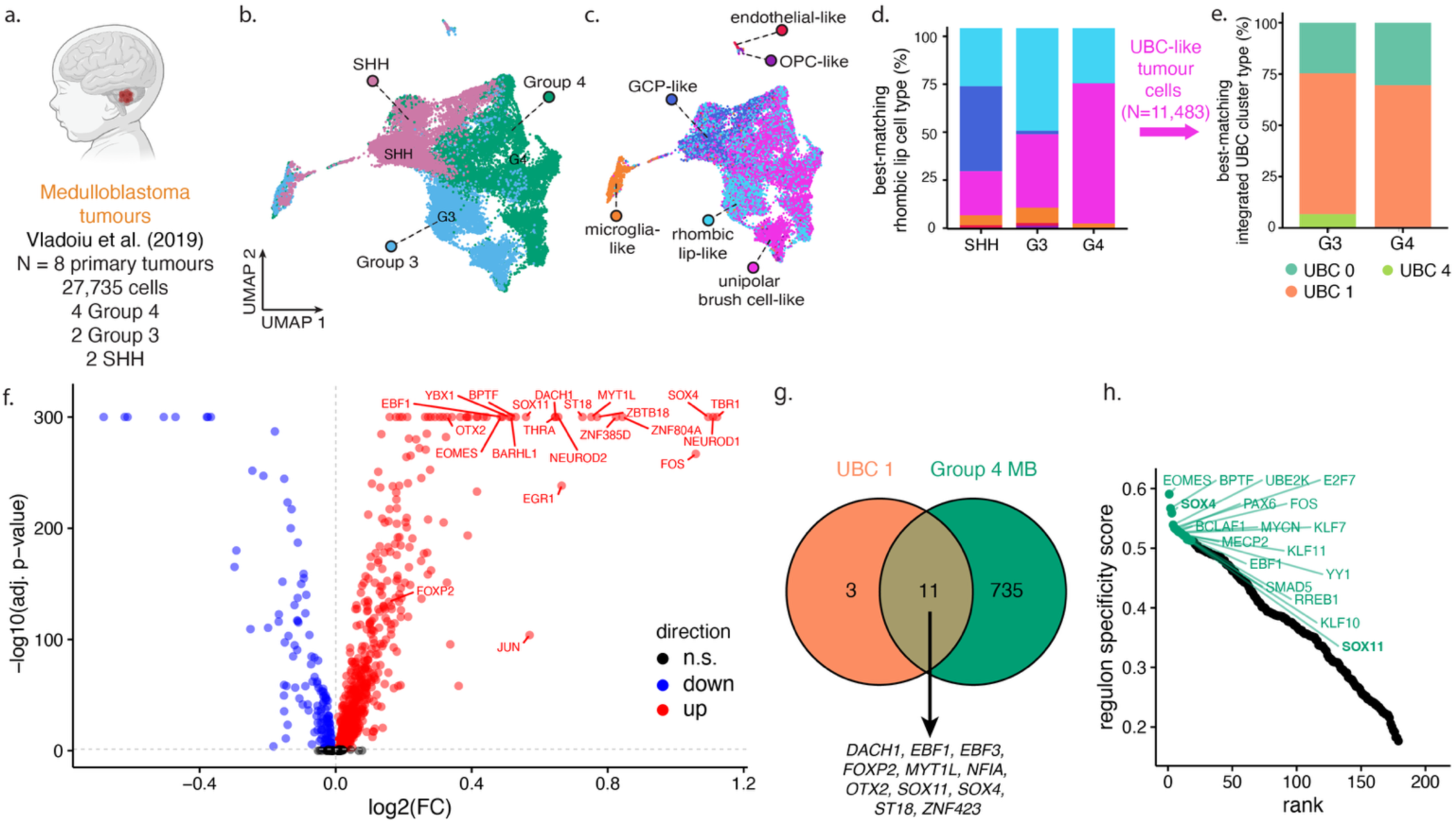
Group 3 and 4 MB tumour cells resemble human-enriched UBC 1 cell state. **a.** Single-cell transcriptomic dataset of MB tumours^23^. **b–c.** UMAP visualization of MB tumour cells, coloured by tumour subtype^23^ (b) and best-matching RL cell type (c). Reference dataset includes non-neuronal cells. **d.** Proportion of MB tumour cells stratified by subtype and best-matching developmental cell type. **e.** Proportion of UBC-like Group 3 and 4 MB tumour cells stratified by subtype and best-matching UBC subtype. UBC subtypes are as those shown in Figure 1e. **f.** Differentially expressed genes in Group 4 MB compared to SHH and Group 3 MB; the top upregulated genes are labelled. Dotted lines indicate a log_2_ fold change (FC) of 0 and an adjusted *p*-value of 0.05. Genes with an adjusted *p*-value of 0 are shown at the top of the plot with their adjusted *p*-value set to 1 × 10^−300^. **g.** Number of transcription factors that are upregulated in Group 4 MB tumour cells and in UBC 1. **h.** Top regulons driving Group 4 MB cell state. GCP, granule cell progenitor; OPC, oligodendrocyte precursor cell; n.s., not significant.

To determine if Group 4 MB tumour cells share drivers of gene expression—*i.e.,* transcription factors (TFs)—with the UBC 1 cell state, we next examined TF expression in these tumour cells. A total of 746 upregulated TFs were identified in Group 4 MB tumour cells (log_2_FC > 0 and Bonferroni-corrected Q < 0.05; Figure 3f). Of these, 11 TFs are also upregulated in the UBC 1 cells, including regulators of neurogenesis such as *SOX4*, *SOX11*, *OTX2*, and *FOXP2* (Figure 3g). We next inferred top regulons in Group 4 MB tumour cells (Figure 3h). The UBC marker *EOMES* was one of the top active regulons for the Group 4 MB cells, consistent with previous research demonstrating the similarity of Group 4 MB tumour cells to UBCs in the developing cerebellum^24^. Similar to UBC 1 cells, *SOX4* and *SOX11* were also among the top regulons for Group 4 MB tumour cells (Figure 3h).

## Discussion

In this work, we identified a population of UBCs that is enriched in humans, arises early in development relative to the parent cerebellar RL progenitor cells, and expresses genes related to development and neurogenesis. This cell population appears to express the retinal photoreceptor gene *EYS*. This population of cells may be driven by the gene regulatory action of SOX4 and SOX11 and transcription factors such as FOXP2. Overlapping top regulons, shared upregulated transcription factors, and the single-cell projection data collectively suggest the possibility that UBC 1 is a cell-of-origin for some subtypes of Group 4 MB tumours.

While the role of *SOX4* and *SOX11* in human cerebellar development has not been studied, both genes have been shown to be important in the expansion and differentiation of progenitor cells in the mouse cerebral cortex^64^. For example, knockout of *Sox11* in mice results in reduced sizes of the cerebellum and the cerebral cortex^65^. Interestingly, *Sox4* activates *Eomes*/*Tbr2*^64^, and a similar mechanism of UBC lineage commitment may be present in the developing human cerebellum. Future work could test the impact of *SOX4* overexpression or downregulation in UBC cell states. Separately, the spatial distribution of the UBC cell states identified here, including the states that are statistically enriched in humans, remains to be resolved. Single molecule fluorescence in situ hybridization (smFISH) in primary tissue from the developing human hindbrain could be performed to verify expression of subcluster-specific markers; for example, *EYS*, *FOXP2*, *CNTNAP2*, and *SOX4* in UBC 1, and *RFX3* in UBC 2.

Using published MB single-cell transcriptomes, we found that *SOX4* and *SOX11* were upregulated in cell clusters of Group 4 MB tumours (Figure 3f, h). This finding is consistent with previous research showing that these genes were overexpressed in primary MB tumours^66^, although this work was published prior to the discovery of the four molecular subgroups of MB, and tumour subtypes were not indicated^20^. Similar to UBC 1 cells, *SOX4* and *SOX11* were found to be among the top regulons of Group 4 MB cells (Figure 3f). Given the role of *SOX4* and *SOX11* in neural proliferation and their high expression in Group 4 MB, we hypothesize that downregulation of these TFs in MB long-passage cell lines will reduce tumour proliferation and promote a more differentiated phenotype. An alternative approach would be to ascertain if overexpression of *SOX4* and *SOX11* in a suitable 2D or 3D human cerebellar neural model^67^ prolongs neuronal maturation. Additional mechanistic studies could be performed to better understand the genes, pathways, and downstream effects *SOX4* and *SOX11* in the context of UBC differentiation and Group 4 MB. Altogether, these results may help identify potential targets for therapy as well as potential diagnostic and prognostic biomarkers. We do not exclude the possibility that other TFs shared between the UBC 1 cell state and Group 4 MB tumours (Figure 3g) may be worthwhile of consideration as candidates to develop laboratory models of Group 4 MB oncogenesis. We also note that the size of the dataset used here (six Group 3 and 4 MB tumours) likely does not capture the heterogeneity of these tumours, and this analysis will need to be repeated in a larger tumour dataset and integrated with genetic and epigenetic tumour data to assess generalizability and develop models of developmental oncogenesis.

An alternative explanation for the presence of the human-enriched UBCs could be temporal and spatial differences in sampling procedures for the source human and mouse datasets used in this work, particularly the human dataset. Given the much shorter timescale of development in mice compared to humans, there may be transient cell states in mouse cerebellar development that may not be present in the mouse datasets due to the lack of sampled timepoints. Integration of gene signatures from UBC 1 and UBC 2 cells with spatial assays (FISH, spatial transcriptomics) of the developing human cerebellum may identify distinct spatial distributions of the UBC clusters, in turn suggesting different models of UBC lineage specification, and disambiguating between EOMES+ UBCs and deep cerebellar nuclei^68^. Separately, UBCs are not uniquely found in the cerebellum, as a population of UBCs also exists in the dorsal cochlear nucleus of the brainstem^7^. As the nuclei for the human datasets used in our study were extracted from cerebellar fragments, contamination from other regions of the hindbrain is unlikely.

A technical consideration arises from the choice of harmonization algorithm for the human-mouse datasets. While the integration shown here was performed using canonical correlation analysis (CCA) in Seurat^31^, integration was also performed using reciprocal principal component analysis (RPCA)^69^ and Harmony^32^. In general, integration with RPCA did not appear to give different results from CCA, and similar cell type clusters were identified (Supplementary Figure 3). However, integration with Harmony appeared to mix cells too strongly, with related but distinct cell types clustering together (Supplementary Figure 3e–f). Indeed, benchmarking of various methods for integrating cells across different species has shown that Seurat CCA and RPCA are among the top performers in terms of biological conservation and species mixing^70^.

A different methodological consideration stems from our use of orthologous genes. Here, we integrated human and mouse cells using genes with one-to-one orthologs. However, evolution has given rise to duplicated and novel genes in humans which may contribute to human-specific features of neurodevelopment^71,72^. The non-orthologous genes excluded from our integration were significantly enriched for 345 pathways that were related to cell division processes, immune system processes, drug metabolism, and chemical perception and response (Q < 0.05; Supplementary Figure 13). Given that this study focuses on neurodevelopment, future analyses could benefit from exploring the role of the non-orthologous human genes, particularly those controlling cell proliferation. While some orthologs have been shown to have human-specific expression compared to other mammals^73,74^ and human-specific cell states have been identified using orthologs^27^, human genes with no known mouse orthologs may also play a factor in driving the unique expansion of the human cerebellum^71^. As such, whether any non-orthologous genes play a role in promoting expansion and proliferation of the human cerebellum will need to be investigated further.

In summary, our analysis revealed two subpopulations of UBCs that are enriched in the developing cerebellum of humans compared to mice. Group 3 and 4 MB tumour cells appear to share multiple features with one of these, an early UBC state here called UBC 1. One direction for future work is the molecular characterization and delineation of the UBC cell lineage in humans and mice that would refine the findings here using complementary approaches such as *in situ* hybridization or immunohistochemistry in primary tissue. Another approach is to use the findings here to validate which TFs have a joint effect on the developing cerebellum as well as support proliferation of MB tumours, using genetic perturbation in preclinical MB models, or 2D or 3D models of the developing cerebellum^67^. Such TFs may be useful in developing novel preclinical models for Group 3 and 4 MB to promote rational therapy design.

## Acknowledgements

BioRender was used to generate some of the figures in this work.

## Data and Code Availability

The integrated human and mouse cerebellar developmental single-cell atlases will be made available via a Zenodo repository. Software to reproduce the analysis in this manuscript will be made available via a GitHub repository under a Creative Commons Attributions License upon publication.

## Supplementary Figures

**Supplementary Figure 1.**
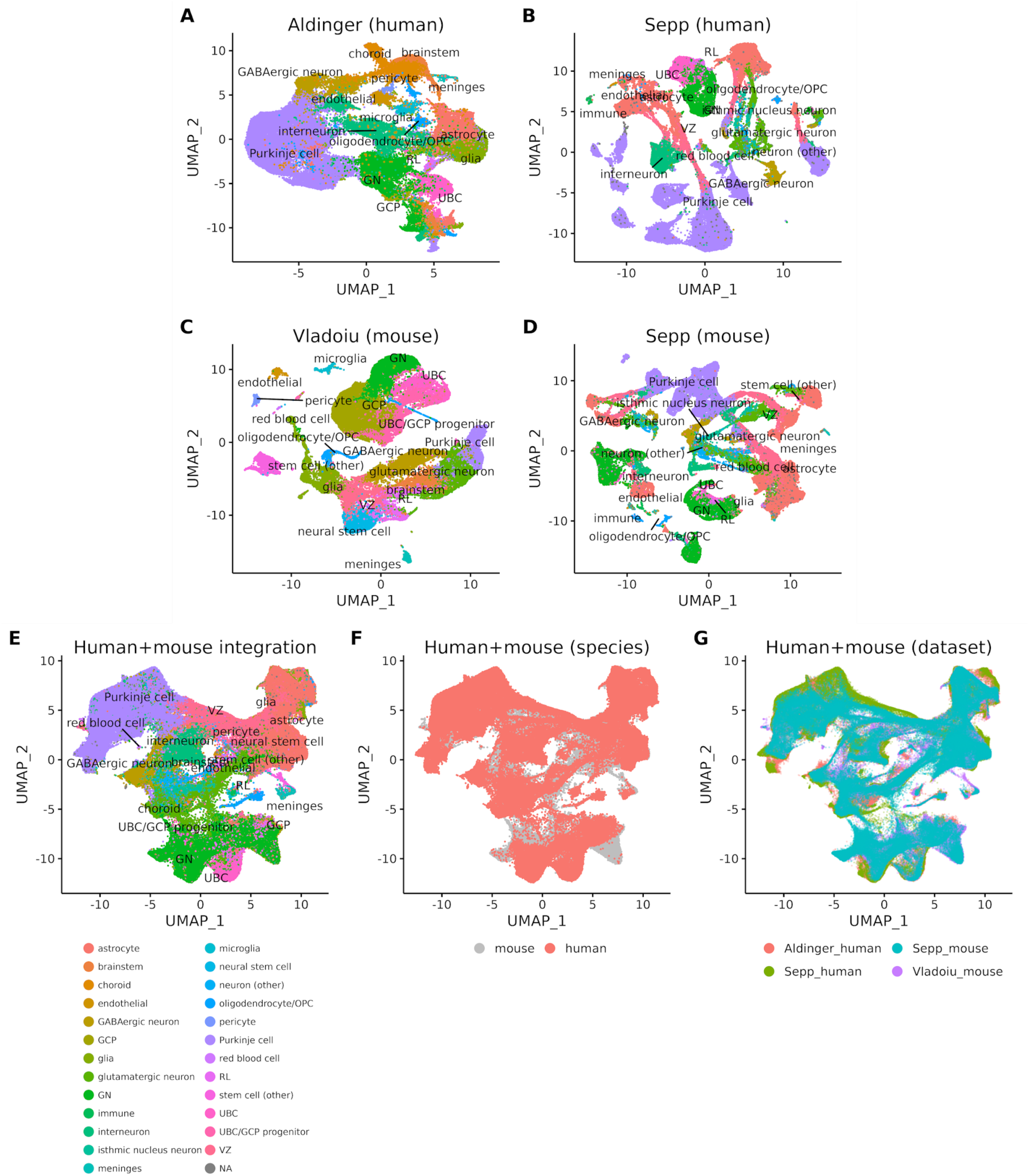
Individual and integrated human and mouse scRNA-seq datasets of the developing cerebellum. UMAP visualizations of each individual dataset (A–D) and following the integration of cells from all four datasets (E–G). Cells are coloured by consolidated cell types based on the annotations from the original datasets (A–E), by species (F), or by dataset (G).

**Supplementary Figure 2.**
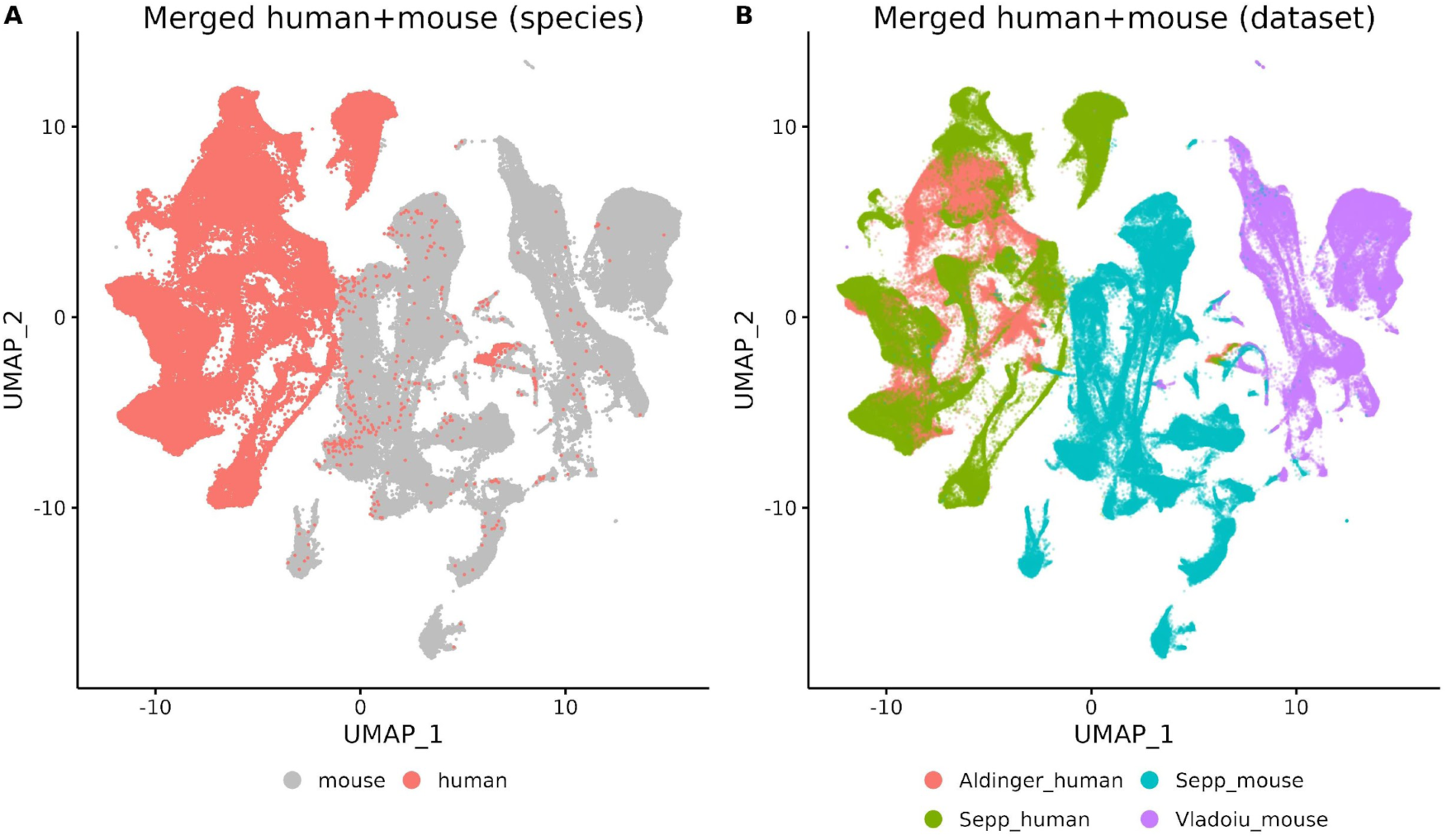
UMAP visualizations of the merged human and mouse cerebellar datasets without integration. Cells are coloured by (A) species and (B) dataset.

**Supplementary Figure 3.**
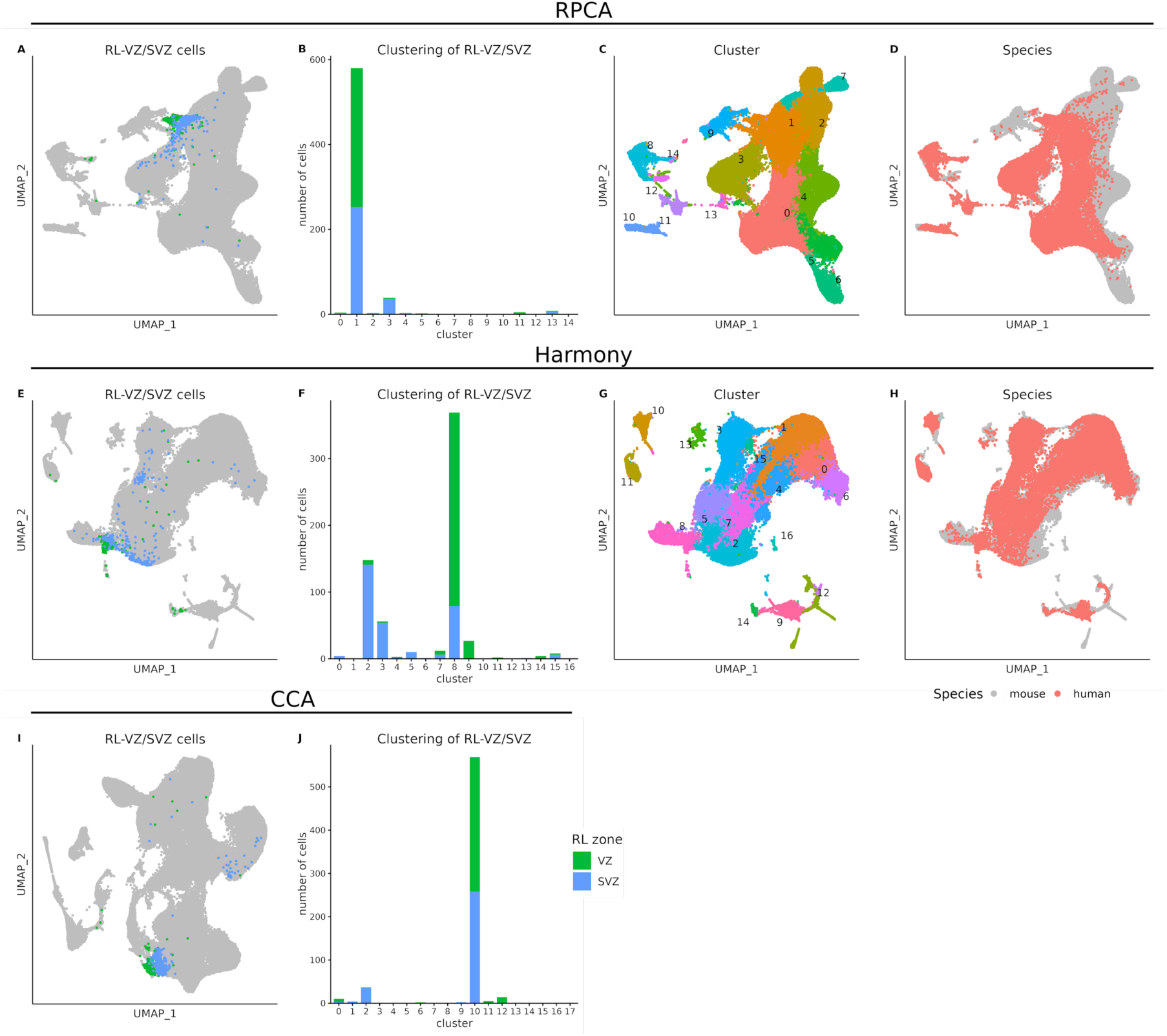
RPCA and Harmony integration of RL lineage cells. (A–D) Integration of the RL lineage cells using RPCA in Seurat. (A) UMAP visualization of the RL-VZ and RL-SVZ cells from the Aldinger dataset^12^ as annotated by Hendrikse *et al.*^24^. (B) Number of RL-VZ and RL-SVZ cells in each cluster. (C–D) UMAP visualizations of RL lineage cells coloured by cluster (C) and species (D). (E–H) Same as (A–D), but for integration with Harmony^32^. (I–J) Same as (A–B), but for integration with CCA. CCA, canonical correlation analysis; RPCA reciprocal principal component analysis; RL, rhombic lip.

**Supplementary Figure 4.**
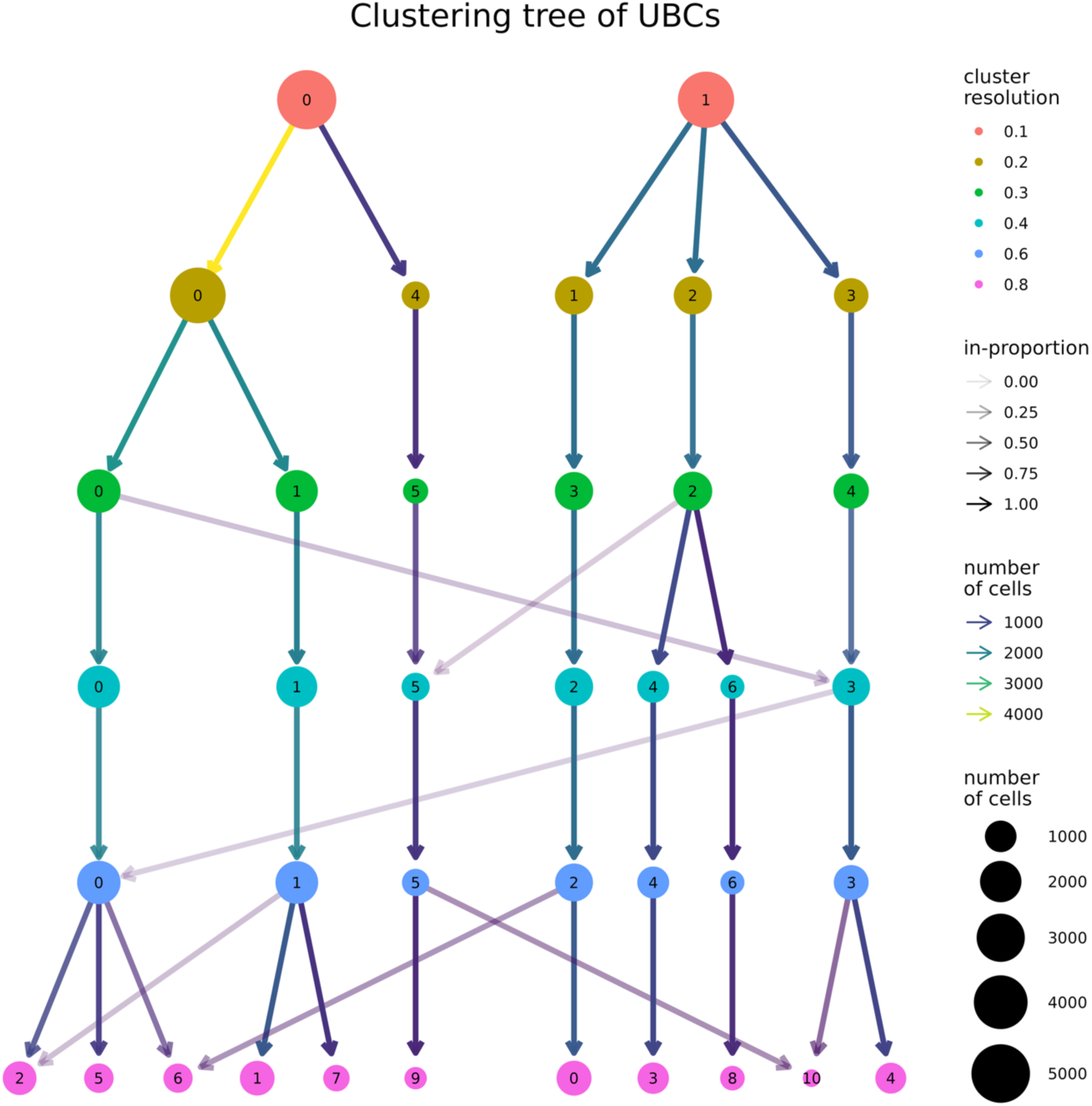
Clustering tree plot of UBCs at different clustering resolutions. This plot shows changes in single-cell cluster membership for integrated human and mouse UBCs across different clustering resolutions. Each node represents a UBC cluster comprised of integrated human and mouse UBCs, and each row shows the clusters generated at a specific resolution from 0.1 to 0.8 (as indicated by the node colour; granularity increases with resolution size). The size of the node represents the number of cells in that cluster. The colour of the arrows represents the number of cells that split from each parent node to the child nodes after the clustering resolution is changed. The opacity of the arrows represents the “in-proportion” metric, which is defined as the ratio of the number of cells in the arrow to the number of cells in the cluster it points toward. The plot was generated using the *clustree* package^33^. UBC, unipolar brush cell.

**Supplementary Figure 5.**
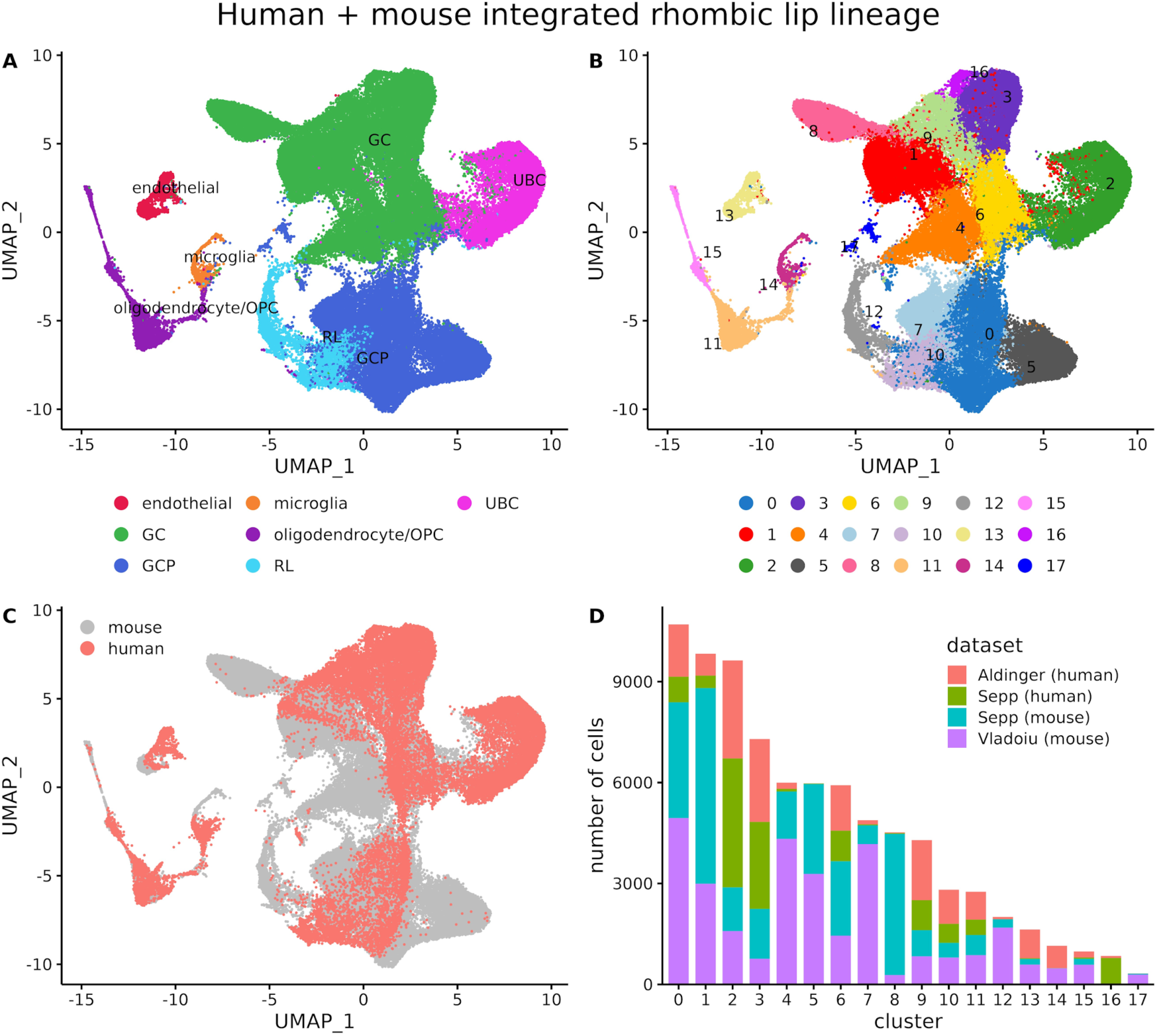
Integration of the cerebellar RL lineage cells. (A–C) UMAP visualization of the CCA-integrated RL lineage cells coloured by (A) cell type, (B) cluster, and (C) species. (D) Bar plot showing the distribution of cells from each dataset in each of the clusters. Panels (A) and (C) are identical to those shown in Figure 1.

**Supplementary Figure 6.**
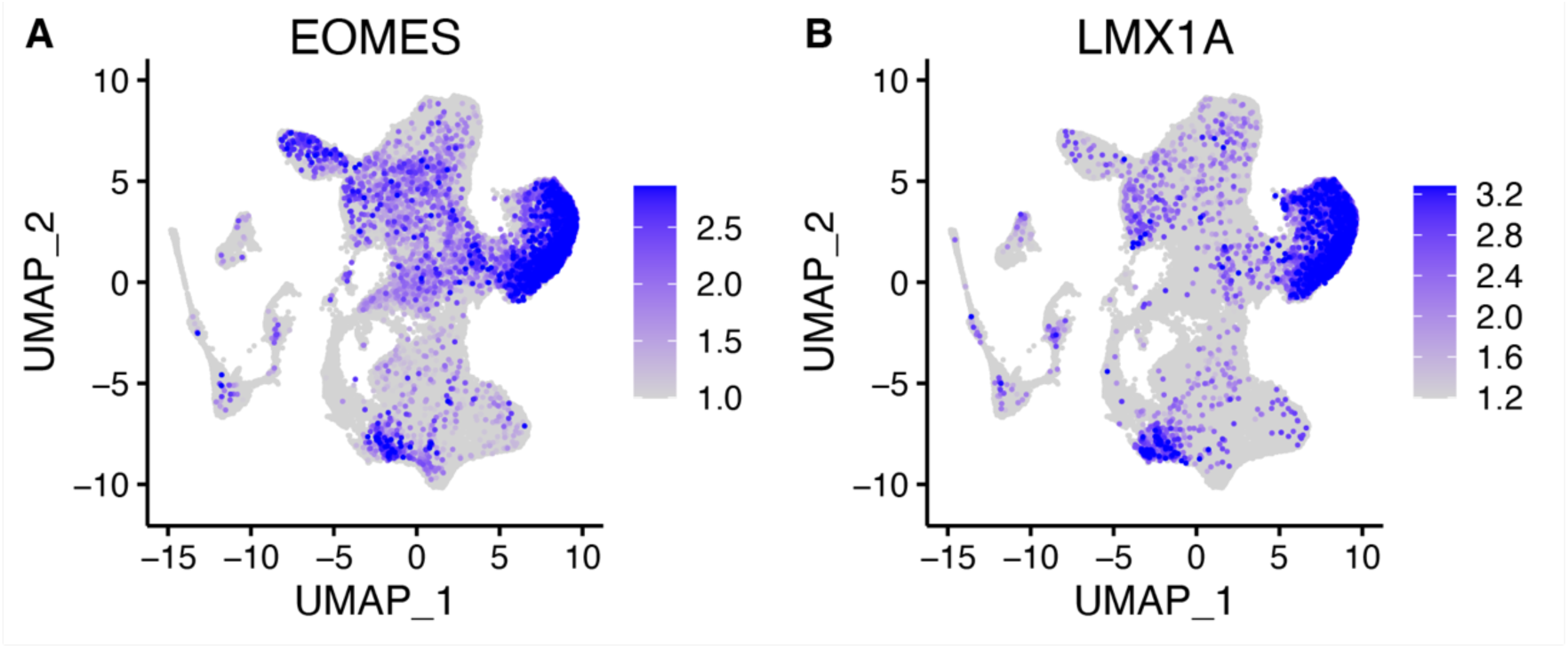
UMAP visualizations showing the expression of the UBC markers *EOMES* (A) and *LMX1A* (B) in cerebellar RL lineage cells.

**Supplementary Figure 7.**
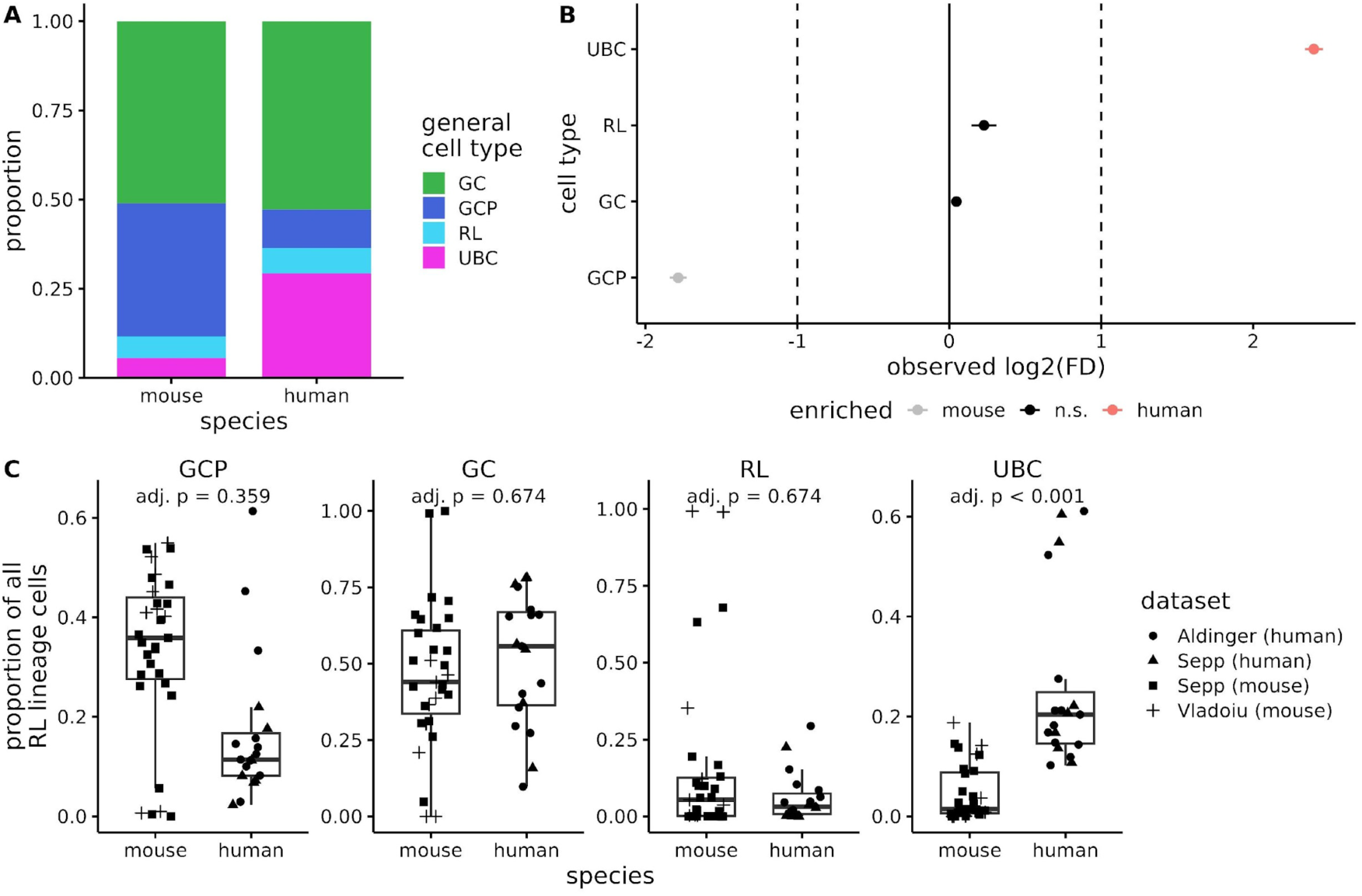
Cell type proportions of RL lineage cells in mice and humans. (A) Bar plot showing the proportion of RL cells, GCPs, GCs, and UBCs in mice and humans. (B–C) Tests for differences in cell type abundance using *scProportionTest* (B) and *propeller* (C). Both show a significantly higher proportion of UBCs in humans compared to mice. (B) *scProportionTest* performs a permutation test (*n* = 10,000 permutations), with the lines showing the 95% confidence interval. Cell types with a log_2_FD > 1 are enriched in humans, and vice versa for cell types with log_2_FD < −1. (C) Box plots showing differences in sample proportions between mice and humans. Each dot represents a biological sample from the dataset. Sample-level information was not available for the Vladoiu dataset, so cells were grouped by age instead. FD, fold difference; n.s., not significant; GC, granule cell; GCP, granule cell progenitor; RL, rhombic lip; UBC, unipolar brush cell.

**Supplementary Figure 8.**
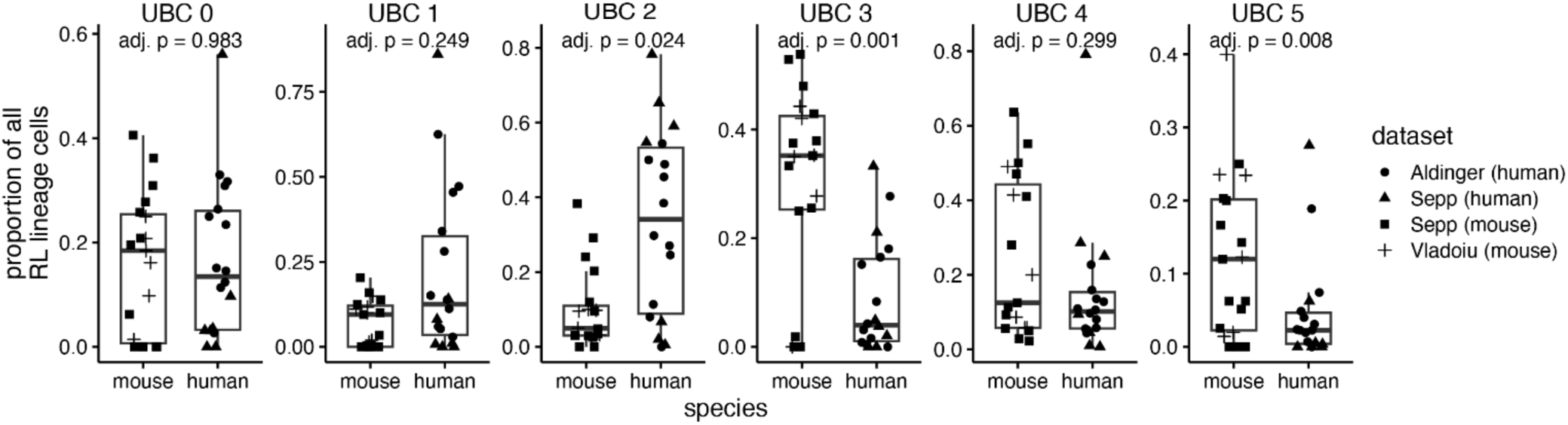
Cell type proportions of UBC subpopulations in mice and humans. Box plots showing differences in sample proportions between mice and humans. Each dot represents a biological sample from the dataset. Sample-level information was not available for the Vladoiu dataset, so cells were grouped by age instead. FD, fold difference; n.s., not significant; UBC, unipolar brush cell.

**Supplementary Figure 9.**
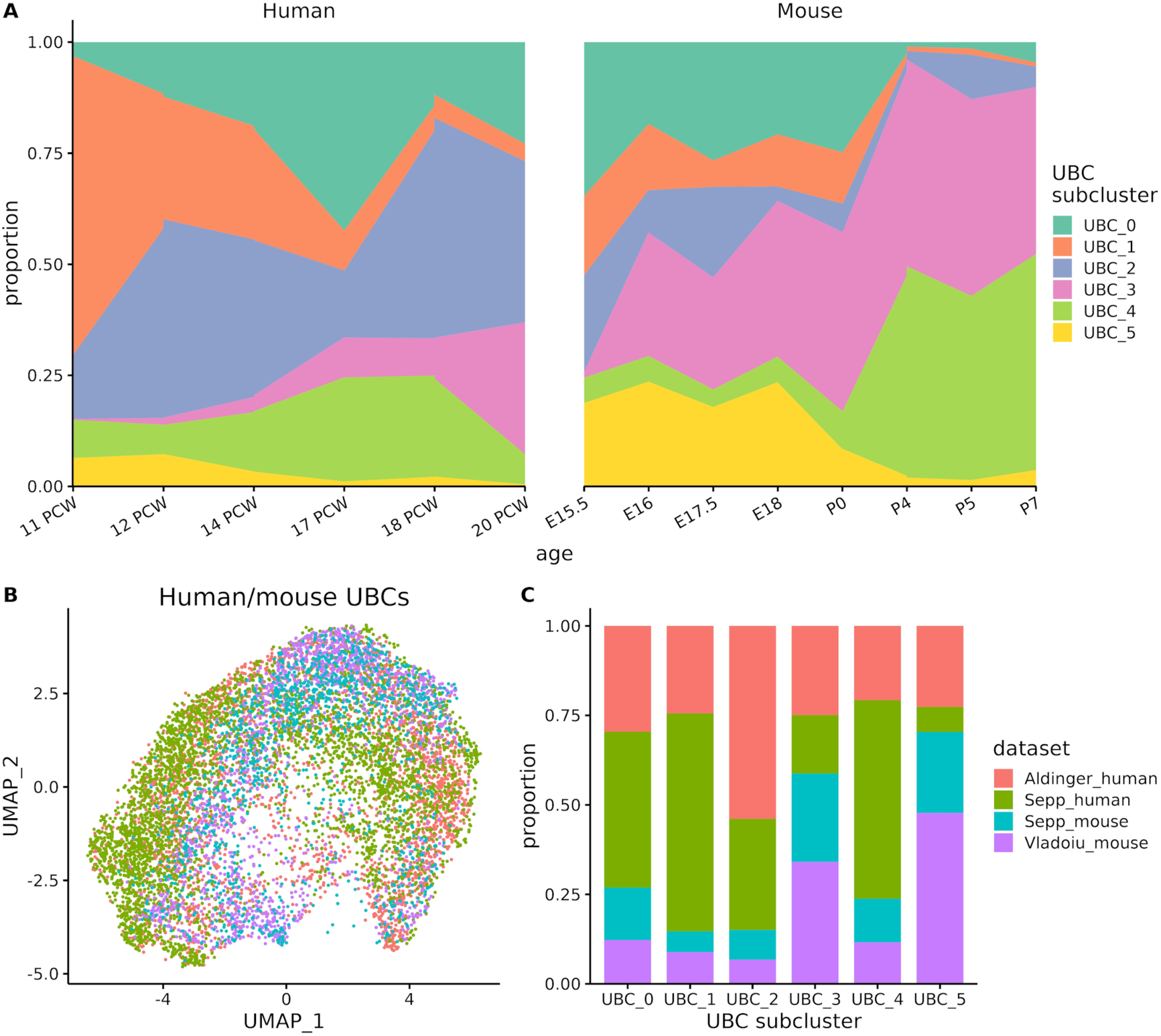
Distribution of ages and datasets in UBC subpopulations. (A) Proportion of human (left) and mouse (right) UBCs in each cluster by age. Ages with fewer than 100 human cells (16 and 21 PCW) or 50 mouse cells (E10–E15 and P14) are not shown. (B) UMAP visualization of the integrated human and mouse UBCs coloured by the original dataset. (C) Proportion of cells from each of the original datasets. Aldinger_human, N = 2,915 cells; Sepp_human, N = 3,832 cells; Sepp_mouse, N = 1,297 cells; Vladoiu_mouse, N = 1,586 cells.

**Supplementary Figure 10.**
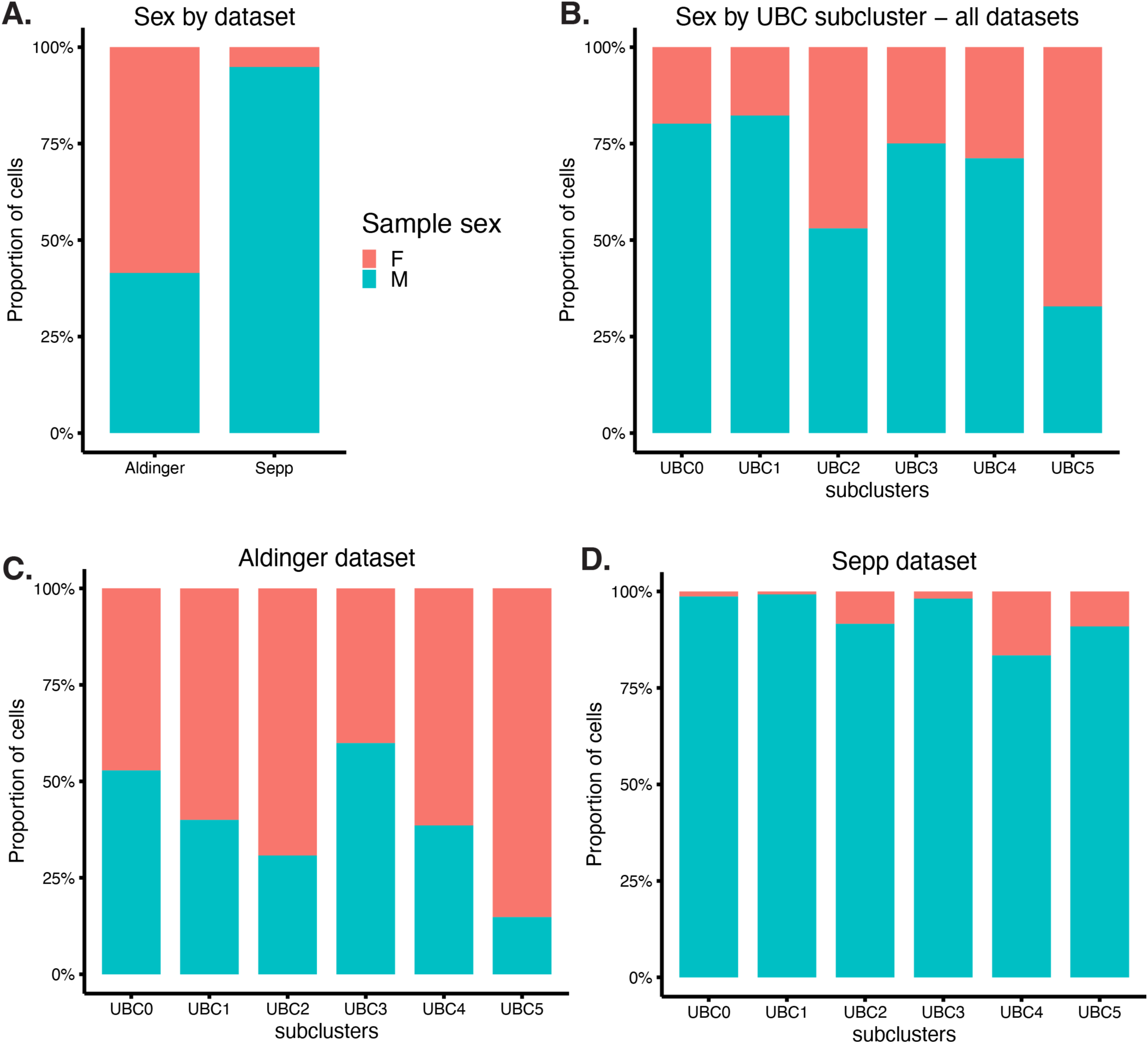
Distribution of donor sex in UBC subpopulations and by dataset. (A) All human unipolar brush cells used in the subclustering analysis, shown by dataset and sex (N = 6,747 human cells in total). (B) Distribution of donor sex by UBC subpopulations. Data shown is for the Aldinger and Sepp datasets combined (same data as in panel A). (C–D) Distribution of donor sex by UBC subpopulations, shown for the (C) Aldinger dataset, and the (D) Sepp dataset.

**Supplementary Figure 11.**
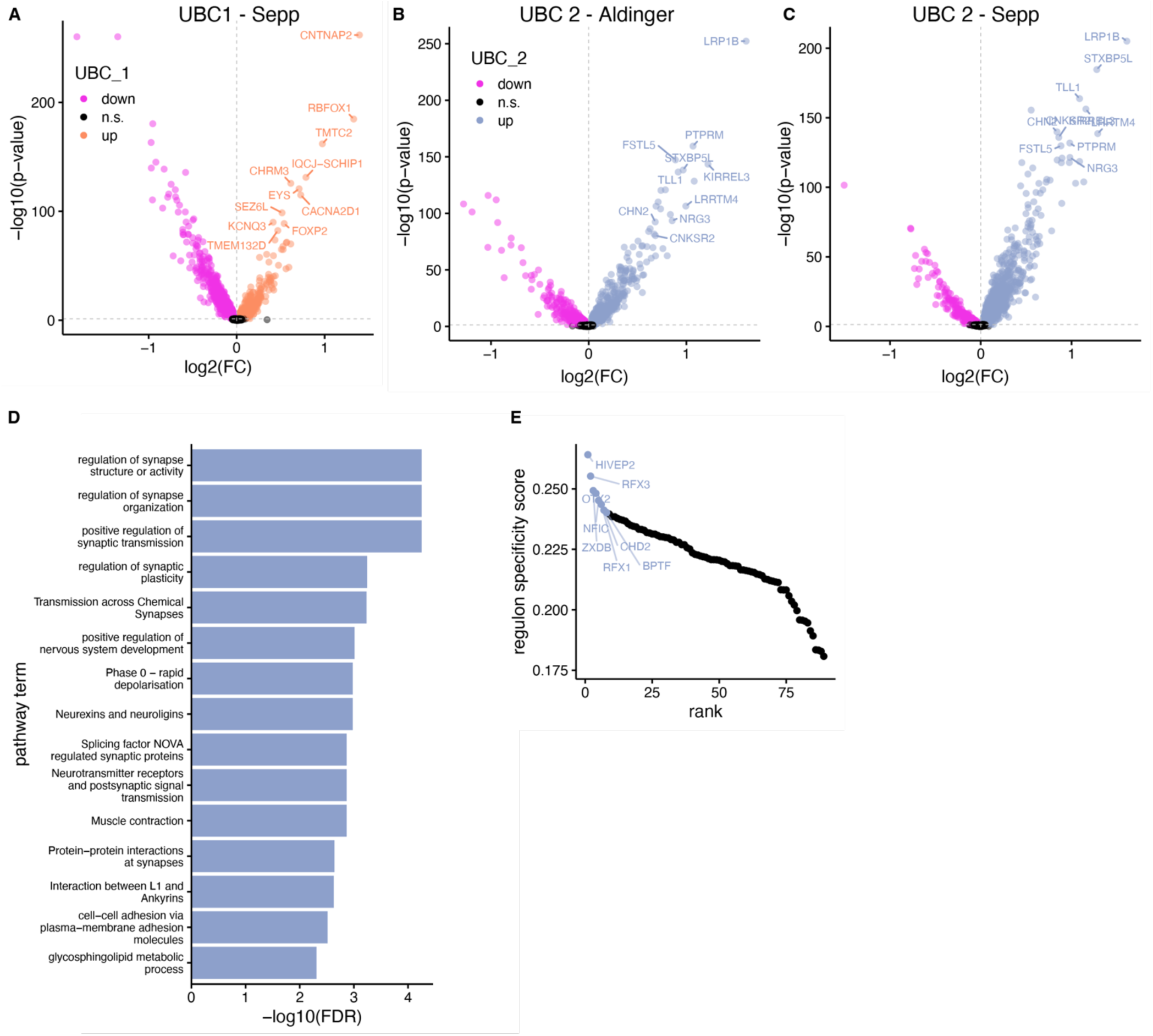
Upregulated genes, pathways, and regulons in UBC 1 and UBC 2 (A). Volcano plot and upregulated genes in the UBC 1 cluster, when only cells from the Sepp cluster are considered (full list of differentially expressed genes in Supplementary Table 3). (B–C) Volcano plots for the UBC 2 cluster, when subset for cells from the Aldinger dataset (B) or cells from the Sepp dataset (C). (D) Top ten enriched pathways for UBC 2 cell cluster; full set of pathways in Supplementary Table 6. (E) Top regulons for UBC 2 cell cluster; full set of regulon specificity scores in Supplementary Table 5.

**Supplementary Figure 12.**
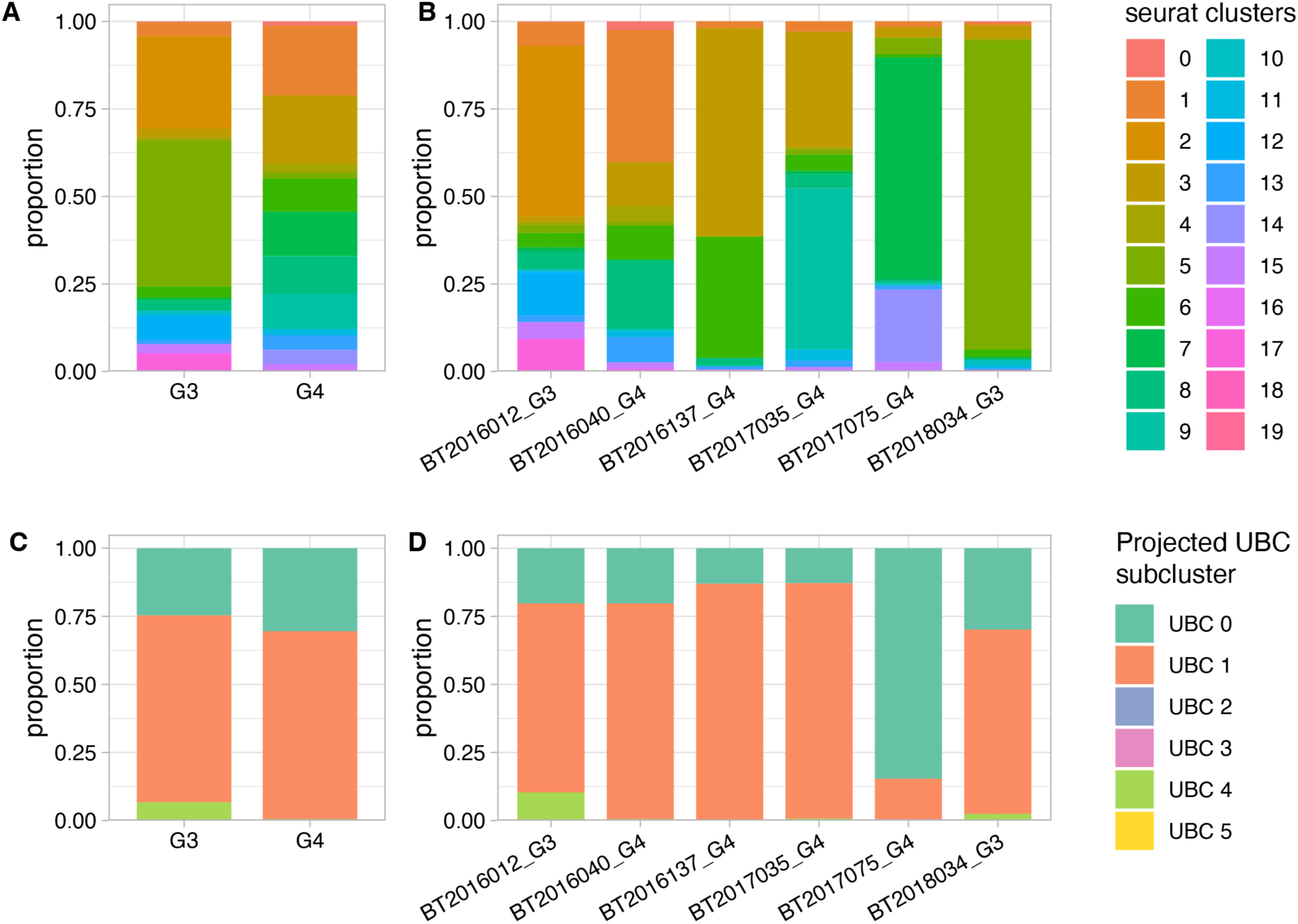
Clusters of UBC-like tumour cells and best-matching UBC subcluster. (A–B) Distribution of UBC-like Group 3 and 4 MB cells according to clusters identified by Seurat. Cluster assignments were made on the full set of 27,735 tumour cells. (A) shows data by tumour subtype, and (B), by tumour sample. (C–D) shows best-matching UBC subcluster as determined by SingleR projections onto UBC-like tumour cells. (C) shows breakdown by subtype and (D) shows breakdown by individual tumour sample.

**Supplementary Figure 13.**
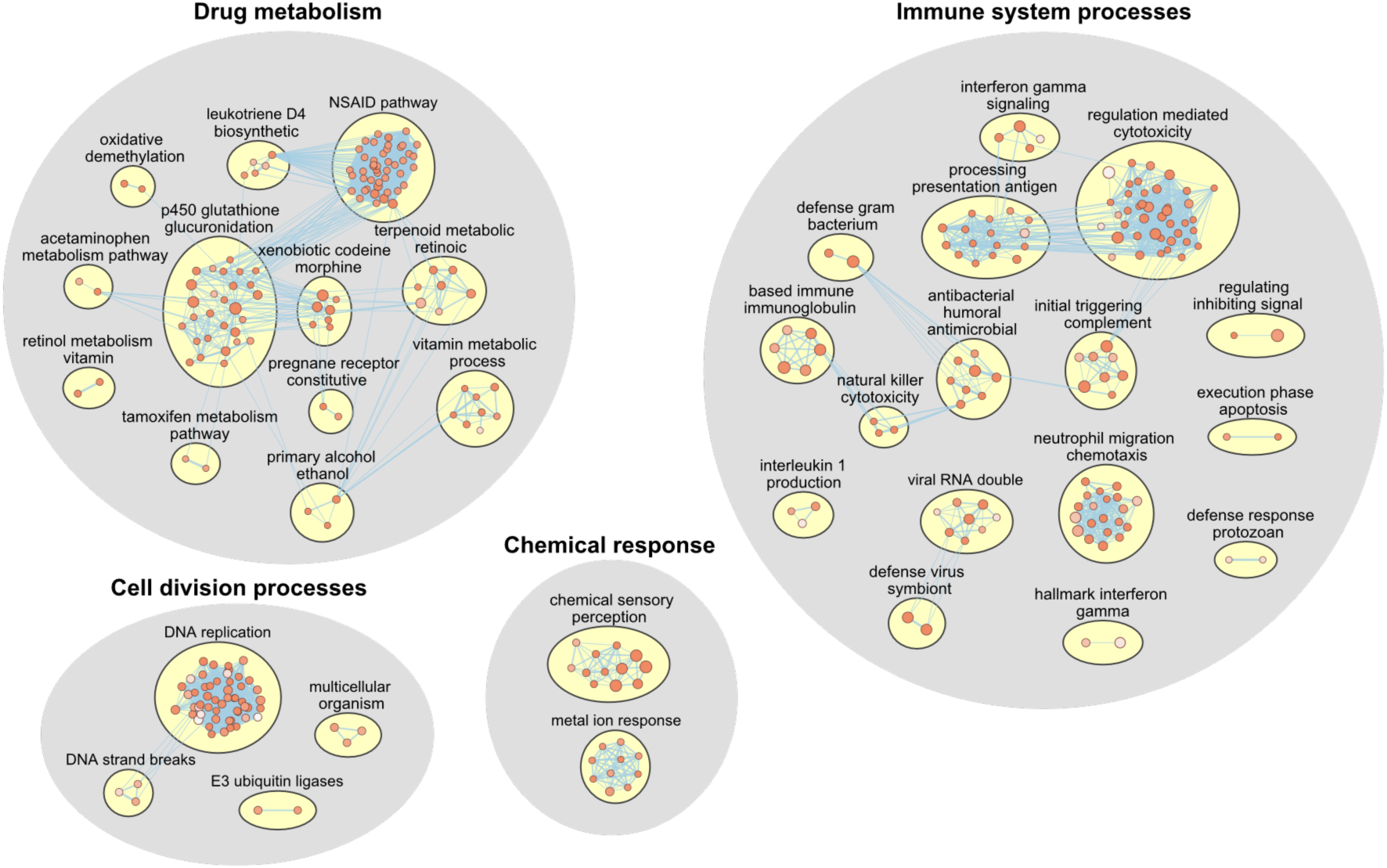
Enriched pathways in non-orthologous human genes. Pathway enrichment analysis on human genes from both human cerebellar datasets^5,27^ that did not have a one-to-one mouse ortholog (non-orthologous genes). A custom Enrichment Map gene set file from the Bader lab was used (see Methods). Each node represents a pathway, and a total of 345 significantly enriched pathways were identified (FDR < 0.05). The enriched pathways were clustered, annotated, and visualized using Enrichment Map and AutoAnnotate in Cytoscape (20 pathways with no overlapping genes not shown here). FDR, Benjamini-Hochberg false discovery rate.

